# Endless Forms Most Beau-toe-ful: Evolution of the Human Hallux

**DOI:** 10.1101/019653

**Authors:** Zachary Throckmorton

## Abstract

**Background:** The adduction of the first pedal ray in humans, such that the hallux is incapable of functional opposability, is a major feature of the evolution of the hominin foot (e.g. Darwin 1872, Haeckel 1879, Latimer and Lovejoy 1990). While hallucal adduction facilitates obligate bipedalism, it inhibits but does not eliminate arboreal grasping ability. The angle of the hallux (as determined by the longitudinal axis of the first metatarsal) relative to the rest of the foot (as determined by the longitudinal axis of the second metatarsal) is a product of both hard-and soft-tissues. Given the failure of soft tissue to fossilize, direct evidence for hallucal angle evolution is scanty, and consensus has not emerged as to when and how the modern human condition of hallucal non-opposability evolved.

**Methodology/Principal Findings:** Analysis of a large sample (n = 331) of radiographs taken from the dorsal aspect of living human feet elucidates the relationship between osteological measures and the magnitude of hallucal adduction, which is resultant of both hard-and soft-tissue anatomy (Lovejoy *et al.* 2009). I describe the correlation of hallucal convergence with first metatarsal posterior articular facet morphology, which allows inference of hallucal convergence in the absence of the medial cuneiform. I report parameters of modern human hallucal convergence variation and offer insight into the hominin fossil record. I infer that the hallucal convergence of the recently reported specimen from Burtele (BRT-VP-2/73) falls within the range of living human variation, inconsistent with the interpretation that this hominin retained the ability to oppose its great toe for the purpose of arboreal locomotion (Haile-Selassie *et al*. 2012). Of the fossil hominin first metatarsals surveyed, all fall within the range of living human variation, consistent with previous research on medial cuneiform (McHenry and Jones 2006) and first metatarsal (Berillon 1999) morphology.

**Conclusions/Significance:** This study finds that the modern human condition of hallucal non-opposability was present in the genus *Australopithecus*. However, while the Burtele specimen (BRT-VP-2/73) falls within the range of living human variation, it displays a markedly divergent hallux compared to most living humans. This study suggests that, as in humans today, there was variation in hallucal divergence in Plio-Pleistocene hominins. Rather than using the terms ‘opposability’ and the ambiguously synonymous ‘grasping,’ I suggest here the term ‘clamping’ (as in the motion performed by a clamp) be used to describe the less powerful and less dexterous ability of modern humans to place and hold a thin, narrow object between the great toe and second toe ray in a limited, albeit functional, manner.

## Introduction

The functional non-opposability of the human hallux is unique among the hominoidea (Cartmill 1985). Hallucal adduction facilitates obligate terrestrial bipedalism. Hallucal adduction is achieved in humans by both hard-and soft tissue differences compared to the apes. While all primates possess a transverse arch, the first plantar and dorsal tarsometarsal ligaments, the first plantar and dorsal intermetatarsal ligaments, the superficial and deep transverse metatarsal ligaments, and the first interphalangeal band of the plantar aponeurosis are in humans reduced in length compared to the hominoidea. These ligamentous interconnections serve to increase the strength, durability, and general patency of not only the transverse arch, but also the medial longitudinal arch. In addition to the adductive tightening of the ligaments of the first pedal ray, its bony elements (first metatarsal, proximal and distal first pedal phalanges) are reduced in length. The shortening of these pedal skeletal elements serves multiple purposes; of particular note, bringing the heads of all metatarsals into alignment produces a more cohesive transverse arch and provides better stabilization of the foot through much of the gait cycle (especially during single limb support, which occurs soon after heel strike, a reduced first ray length facilitates better locking of both the forefoot). Concomitant with the shortening of these bones, the *m. flexor hallucis longus*, *m. tibialis anticus*, and *m. fibularis longus* tendons, as well as the *m. flexor hallucis brevis* and *m. adductor hallucis* are reduced in length compared to the apes. Decreasing the lengths of these muscles and their tendinous insertions affords a much more stable medial longitudinal column. Lastly, the morphology of the medial tarsometatarsal joint further contributes to the adduction of the hallux. The obliquity of both the articular facet of the medial cuneiform with the first metatarsal and the articular facet of the first metatarsal with the medial cuneiform (the medial tarsometatarsal joint) in humans is reduced compared to apes (Berillon 1999); this configuration limits mobility but greatly increases stability.

Given the anatomical complexity of the first pedal ray’s relationship to the foot, consensus has not emerged as to how its adduction evolved in the hominin lineage. This anatomical complexity provides myriad targets for natural selection and genetic drift to operate on the hallucal complex. Not only the paucity of the fossil record, but also the failure of relevant soft tissues to fossilize, have impeded understanding of the pattern and timing of this important feature’s evolution. This lack of clarity has led to variable interpretations of fossil foot bone anatomies by researchers. Early hominins are most relevant to this discussion as *Homo* is broadly assumed to look modern.

Though the proximal base of the *Ardipithecus ramidus* first metatarsal (ARA-VP-6/500-089) is abraded and fragmentary (Lovejoy *et al.* 2009) (and therefore not considered in this study), the first tarsometarsal articular facet of the medial cuneiform (ARA-VP-6/500-088) is well-preserved. Lovejoy *et al.* (2009) illustrate that its hallucal abducence – largely inferred from the morphology of the medial cuneiform - is outside of the range of modern humans and comfortably within the ranges of *Pan* and *Gorilla*. That is, *Ar. ramidus* exhibited complete opposability of its great toe as in extant African apes; Lovejoy *et al*. (2009) argue it is the only known hominin to exhibit this condition.

Multiple first metatarsals assigned to *Australopithecus afarensis* have been recovered from the Hadar locality of Ethiopia (Latimer *et al*. 1982). Both A.L. 333-115 and A.L. 333-21 preserve only the distal aspect and are not considered in this study. A.L. 333-54 presents the proximal half of the bone, and its proximal articular facet is well preserved. Latimer and Lovejoy (1990) interpret the first tarsometatarsal joint as quite human-like, with both the morphology of the distal articular facet of the medial cuneiform and proximal articular facet of the first metatarsal suggesting “total absence of significant hallucal mobility.” However, Proctor *et al.* (2008), utilizing quantitative three-dimensional shape analysis of the proximal articular facets of humans, apes, and fossils, argue A.L. 333-54 exhibits proximal oblique curvature more like apes than humans – though they did not interpret their finding’s potential functional implications. The Laetoli footprints are similarly controversial. Stern and Susman (1983) argue these footprints lack many characteristics typical of modern human-made footprints, such as a medial longitudinal arch indistinguishable from that of extant apes (though see DeSilva and Throckmorton 2010), and the noticeable depression at the head of the first metatarsal caused by a distinct toe-off phase of the gait cycle. Similarly, Tuttle (1981, 1984) argued that while the Laetoli footprints were likely made by an obligate terrestrial biped, the short pedal phalangeal lengths preserved in the ash were incompatible with his interpretation of the lengths of the Hadar fossil feet (i.e. the Hadar hominins did not produce the Laetoli trackways). However, White and Suwa (1987) illustrated how the Laetoli footprints are compatible with *A. afarensis* foot skeletons. That said, neither Stern and Susman (1983) nor did Tuttle (1981, 1984) argue that the Laetoli track makers were ‘transitional’ in their foot anatomy and mode of bipedalism because of a more abducted hallux.

Stw 573 from Member 2 of Sterkfontein, South Africa dating to between 3.0 and 3.5Mya is described as intermediate between modern humans and African apes in its hallucal divergence. While the lateral aspect of the medial cuneiform’s (Stw 573c) tarsometatarsal articular facet appears human-like, its medial aspect appears more ape-like and the proximal articular facet of the first metatarsal is also described as ape-like, “traits associated with a divergent, mobile, opposable hallux” (Clarke and Tobias 1995). However, in their multivariate analyses of the Stw 573 and OH 8 medial cuneiforms employing 11 linear measurements, Kidd and Oxnard (2005) found that both OH 8 and Stw 573 were much more like modern humans than apes. The ca. 1.8Mya SKX 5017 first metatarsal from Member 1 of Swartkrans, South Africa attributed to *Paranthropus robustus* is described as robust, short, with a well-developed proximal diaphysis exhibiting no characters suggesting any degree of functional opposability as in African apes (Susman and Brain 1988). Like A.L. 333-54, Proctor *et al.* (2008) assert SKX 5017 displays more ape-than human-like oblique curvature. Broadly similar to SKX 5017, the SK 1813 first metatarsal from Swartkrans, South Africa (either Member 1 or possibly Member 2) with no taxonomic assignation and of unknown antiquity is robust and short (Susman and de Ruiter 2004) as in modern humans. This specimen has not been explicitly described as evincing ape-like divergence, though both Susman and de Ruiter (2004) and especially Zipfel and Kidd (2006) stress that the proximal articular morphology of neither SKX 5017 nor SK 1813 are clearly more like either African apes or modern humans, at least in respect to these specimens’ proximal articular facet heights and widths.

Furthermore, Susman and de Ruiter (2004) describe the SKX 5017, SK 1813, and Stw 562 (from Member 4 of Sterkfontein, South Africa) first metatarsals as exhibiting head torsion intermediate between that of modern humans and apes, better positioning the hallux for opposition during grasping than in modern humans, but not as well as in chimpanzees.

First metatarsals recovered from specimens assigned to the genus *Homo* display morphology more consistent with the typical modern human pattern in regard to hallucal adduction. Though there is disagreement about the hallucal adduction of *Australopithecus* (Susman *et al.* 1984, White and Suwa 1987, Latimer and Lovejoy 1990), the non-abducent hallux is a clear autapomorphy of *Homo.* Dating to approximately 1.84Mya (Blumenschine *et al.* 2003), Olduvai Hominid 8 preserves much of the first metatarsal, including its proximal base. Initially described as possessing most characters of a derived, human-like foot (Leakey *et al*. 1964) (though Wood 1974 emphasized OH 8’s non-*Homo* characters, these are not immediately relevant to the hallux). The first tarsometatarsal joint surfaces of both the medial cuneiform and first metatarsal are remarkably flat, suggesting minimal, non-functional abducence of the hallux as seen in modern humans (Berillon 1999, Harcourt-Smith and Aiello 1999, McHenry and Jones 2006). First metatarsals recovered from the ca. 1.8Mya site of Dmanisi, Republic of Georgia (D2671 and D3442) are somewhat similar to that of OH 8, SKX 5017, and SK 1813, especially in their relatively small sizes and narrow heads (less like the relatively larger first metatarsals with broader heads typical of modern humans) (Lordkipanidze *et al*. 2007). However, the Dmanisi first metatarsals are remarkably robust (in the upper range of modern variation), display minimal diaphyseal torqueing, and their proximal articular facet morphology suggests that in respect to the divergence of the hallux, they were functionally similar to modern humans in being entirely adducent (Pontzer *et al*. 2010).

The Jinniushan first metatarsal (84.J.A.34), absolutely dated to between 200Kya and 260Kya (Chen *et al*. 1994) and assigned taxonomically to archaic *Homo sapiens* (Bae 2010), does not suggest any degree of functional abduction. Its anatomy is similar to that of many modern humans: its shaft is straight, the metatarsal head is broad and dorsally extended, and the *m. flexor hallucis brevis* medial tendon groove is more pronounced than its lateral tendon groove (Lu *et al*. 2011); its only features that are remarkable compared to a typical modern human first metatarsal are its generalized robusticity and the plantar ‘beaking’ of its head. Similar to the Jinniushan first metatarsal, the first metatarsals assigned to Neanderthals are, as typical of Neanderthal skeletal elements, robust compared to those of modern humans. Though Boule (1911) and Morton (1926) argued for a degree of Neanderthal hallucal abduction intermediate between modern humans and African apes, their interpretations have not withstood modern scrutiny; Trinkaus (1983) argued that Neanderthal halluces were fully, habitually adducted as they are in modern humans. The pedal remains of *Homo floresiensis* from Liang Bua, Indonesia include a well-preserved left first metatarsal (LB1/21). Its length relative to the associated second, third, fourth, and fifth metatarsals is short compared to most modern humans but falls within the modern human range, it is robustly built, and its proximal articular facet with the medial cuneiform is flat, indicating an adducted non-grasping hallux (Jungers *et al.*2009).

Haile-Selassie *et al*. (2012) describe the BRT-VP-2/73 first metatarsal as mosaic in its character suite. It lacks the nonsubchondral isthmus of its dorsoproximal articular margin found in *Ardipithecus ramidus*, is relatively short compared to the associated second and fourth metatarsals as in African apes, but has a tall hallucal base relative to the shorter bases of the associated metatarsals as in modern humans. They argue on the bases of the short first metatarsal and proximal pedal phalanx as well as medially torqued second metatarsal that BRT-VP-2/73 had an abducent hallux that facilitated grasping capacity and therefore arboreal substrate exploitation more effectively than penecontemporaneous *Australopithecus afarensis* individuals, despite the Burtele specimen’s lack of an associated medial cuneiform.

In this context, I suggest a potential skeletal correlate of hallucal convergence in hominins that requires only the proximal articular facet and diaphysis of the first metatarsal, its proximal articulation angle. Because the relationship of the great toe with the rest of the foot is a product of both soft-and hard tissues, this correlation is most appropriately extracted from living human feet. I test the relationship between the first metatarsal’s proximal articulation angle and the magnitude of hallucal convergence in radiographs of modern human feet (n = 331). The fossil record is then interpreted in light of this analysis to reevaluate ideas concerning the evolution of hominin hallucal adduction.

## Materials and Methods

Approximately 600 dorsal, weight-bearing (during normal standing with the knee in close-packed position) radiographs of the foot taken as part of routine medical care of a modern, habitually shod population of mixed sex and ancestral background were analyzed for inclusion in the study. These x-rays were entirely deidentified prior to analysis. Completely anonymized radiographs are not considered human subjects by the IRB (UW-Madison IRB Exemption M-2010-1265) and research on them is permitted under HIPAA laws. The large sample size facilitated conservative inclusion of normal, skeletally mature individuals’ radiographs. That is, all x-rays that did not allow immediate and unambiguous identification of relevant osteological landmarks due to insufficient resolution (particularly of the first tarsometatarsal joint) were excluded. Radiographs of individuals exhibiting any evidence of trauma, surgical intervention, advanced diabetic neuropathy, Charcot-Marie-Tooth neuropathy, hallux rigidus, hallux valgus, hallux varus, metatarsus primus elevatus, or any other pathological condition that compromises normal foot anatomy and biomechanics were also excluded from the study. This resulted in a sample size of 331 individuals exhibiting variable development of hallucal convergence (Fig. 1). All measurements were taken of the right foot with standard equipment (i.e. viewing box, straightedge, compass). Fossil hominin first metatarsals were photographed dorsally while in anatomical position; this resulted in a sample size of 8 (Table 1).

**Table 1.** Measurements of fossil hominin first metatarsal proximal articulation angles (PAAs) considered in this study.

| PAA (degrees) | Fossil Specimen | Age | Taxonomic Affinity | Reference |
| --- | --- | --- | --- | --- |
| 87 | SKX 5017 | 1.8Mya | *Paranthropus robustus* | Susman and Brain 1988 |
| 88 | 84.J.A.34 | 260-200Kya | Archaic *Homo sapiens* | Lu *et al.* 2011 |
| 90 | OH 8 | 1.8Mya | *Homo habilis* | Leakey *et al.* 1964 |
| 90 | LB1/21 | 18Kya | *Homo floresiensis* | Jungers *et al.* 2009 |
| 95 | SK 1813 | 1.8-1.5Mya | (not assigned) | Susman and de Ruiter 2004 |
| 97 | D3442 | 1.8Mya | *Homo erectus* | Lordkipanidze *et al.* 2007 |
| 98 | Stw 562 | 2.6-2.0Mya | (not assigned) | Susman and de Ruiter 2004 |
| 100 | BRT-VP-2/73 | 3.4Mya | *Australopithecus spp.* | Haile-Selassie *et al.* 2012 |

**Figure 1.**
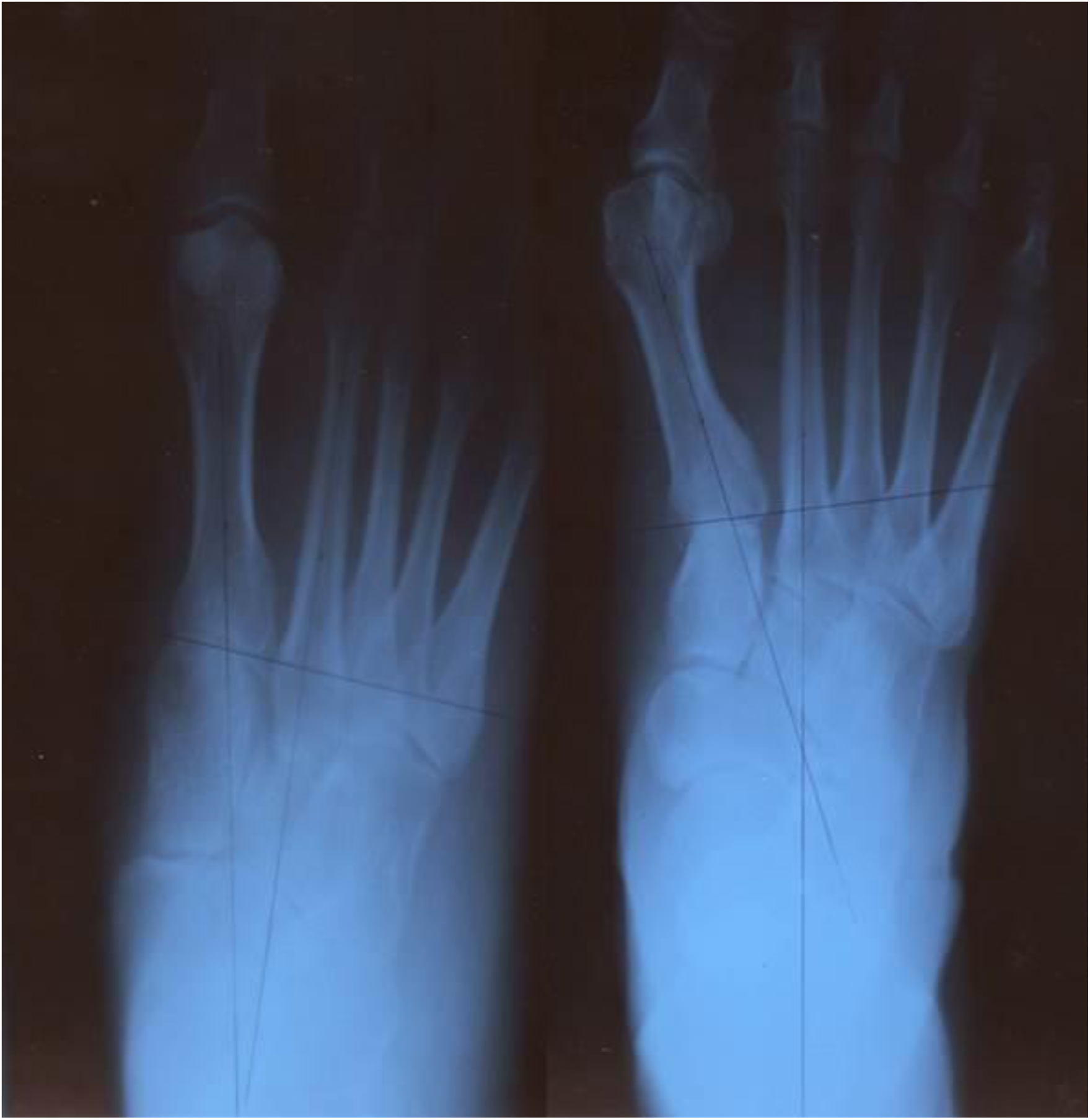
Variation of hallucal convergence in living humans. Two individuals exhibiting a more adducted (convergent) hallux (left) and a more abducted (divergent) hallux (right).

The two measurements collected were the proximal articulation angle of the dorsal aspect of the first metatarsal (proximal articulation angle, PAA) and the angle of the hallux relative to the second ray (great toe angle, GTA) (Fig. 2). The angle of the hallux relative to the second ray (GTA) was determined by drawing lines through the longitudinal axes of the first and second metatarsal diaphyses posteriorly until they intersected (when this intersection occurred posterior to the limit of the actual radiographic film, a piece of standard tracing paper was used to extend the drawing surface to ensure accurate continuity and intersection of the lines). The PAA was determined by drawing a line across the proximal articulation of the first metatarsal’s dorsal aspect compared to the longitudinal axis of the first metatarsal diaphysis (as used in establishing the GTA). A PAA of 90° reflects perpendicularity of the first metatarsal’s longitudinal axis with its proximal articular facet’s dorsal aspect. A PAA of greater than 90° indicates a medially oblique first metatarsal proximal articular facet while a PAA of less than 90° indicates a laterally oblique first metatarsal proximal articular facet. That is, the bony morphology of the first metatarsal’s proximal articular facet itself contributes to an abducted, divergent hallux when its PAA is greater than 90°, and contributes to an adducted, convergent hallux when its PAA is less than 90°. However, the terms abducted, divergent, adducted, and convergent are relative in this context; none of the living humans surveyed in this study have truly abducted, divergent great toes capable of functional opposability as see in extant hominoidea. Based upon comparisons of shod with unshod modern human first metatarsals, Proctor (2010) found no differences between the two groups’ proximal first metatarsal articular facet morphology, increasing confidence in the appropriateness of comparing shod modern human first metatarsals to fossil hominin first metatarsals. Further, Thompson and Zipfel (2005) found no differences between the forefoot widths (determined in part by the degree of hallucal divergence) of shod and unshod humans. However, D’Aout and Aerts (2008) and D’Aout *et al*. (2009) found that habitually unshod feet are wider than habitually shod feet; that is, it is possible that the habitually shod population sampled in this study underestimates the magnitude of hallucal divergence in modern humans. The covariance of the GTA and PAA was uncovered using a Pearson correlation test (Microsoft Excel 2010). To test the probability of sampling PAAs in the fossil hominin first metatarsal from a modern human population, resampling statistics were utilized.

**Figure 2.**
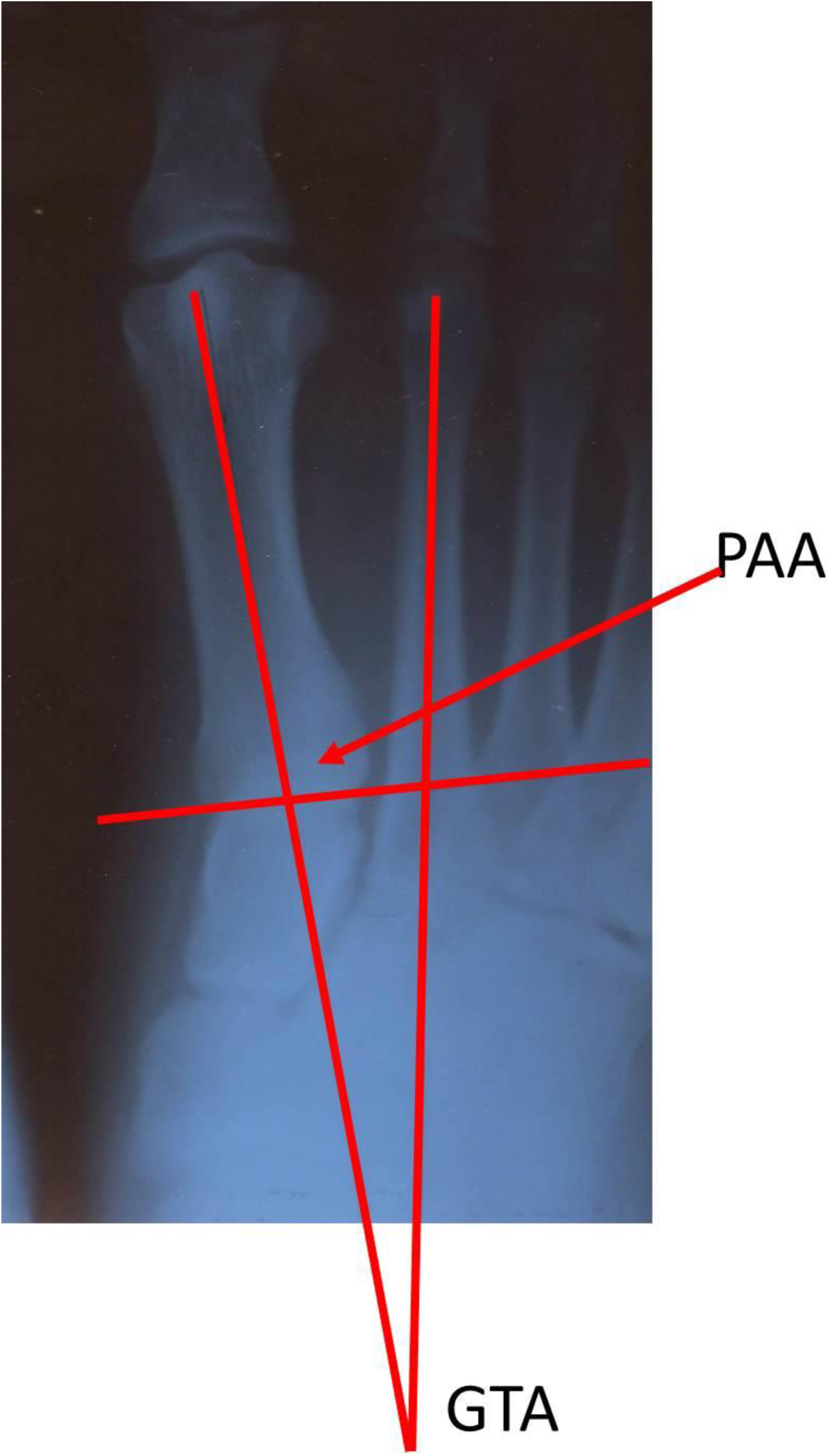
Measurements taken in this study. The first metatarsal proximal angle of articulation (PAA) and the angle of the hallux relative to the foot (GTA).

## Results

To test the hypothesis that the proximal articulation angle (PAA) is an indicator of the great toe angle (GTA), 331 weight-bearing radiographs of human feet taken in dorsal view were examined. A medially oblique PAA is present in 80% of living humans, with 5% and 15% of living humans possessing perpendicular or laterally oblique PAAs, respectively. Parametric statistics reveal PAA is correlated with GTA (r = 0.26, p < 0.001) (Fig. 3). Slightly more than 7% of living humans have a PAA equal or even more medially oblique than BRT-VP-2/73. Thus, the PAA of BRT-VP-2/73 falls within the range of living humans sampled in this study, none of which have functionally opposable great toes. The Woranso-Mille first metatarsal’s PAA is highest and therefore the most extremely medially oblique of fossil hominin first metatarsals surveyed in this study. Stw 562 is slightly less medially obilique than the Burtele specimen. Interestingly, ca. 1.8Mya D3442 from Dmanisi, Republic of Georgia, has the next most medially oblique PAA of the fossil hominins surveyed. Though Stw 573 (more than 3Mya) is incomplete, it appears to be slightly less medially oblique than the Dmanisi specimen. The australopithecine SK 1813 is slightly less medially oblique than D3442. The first metatarsals of the OH 8 foot and the LB1/21 *Homo floresiensis* foot have a perpendicular PAA, while the specimens 84.J.A.34 from Jinniushan, China and SKX 5017 from Swartkrans, South Africa actually have slightly laterally oblique PAAs – though in the case of 84.J.A.34, slight abrasion of the proximal articular facet laterally precludes great confidence in its PAA measurement.

**Figure 3.**
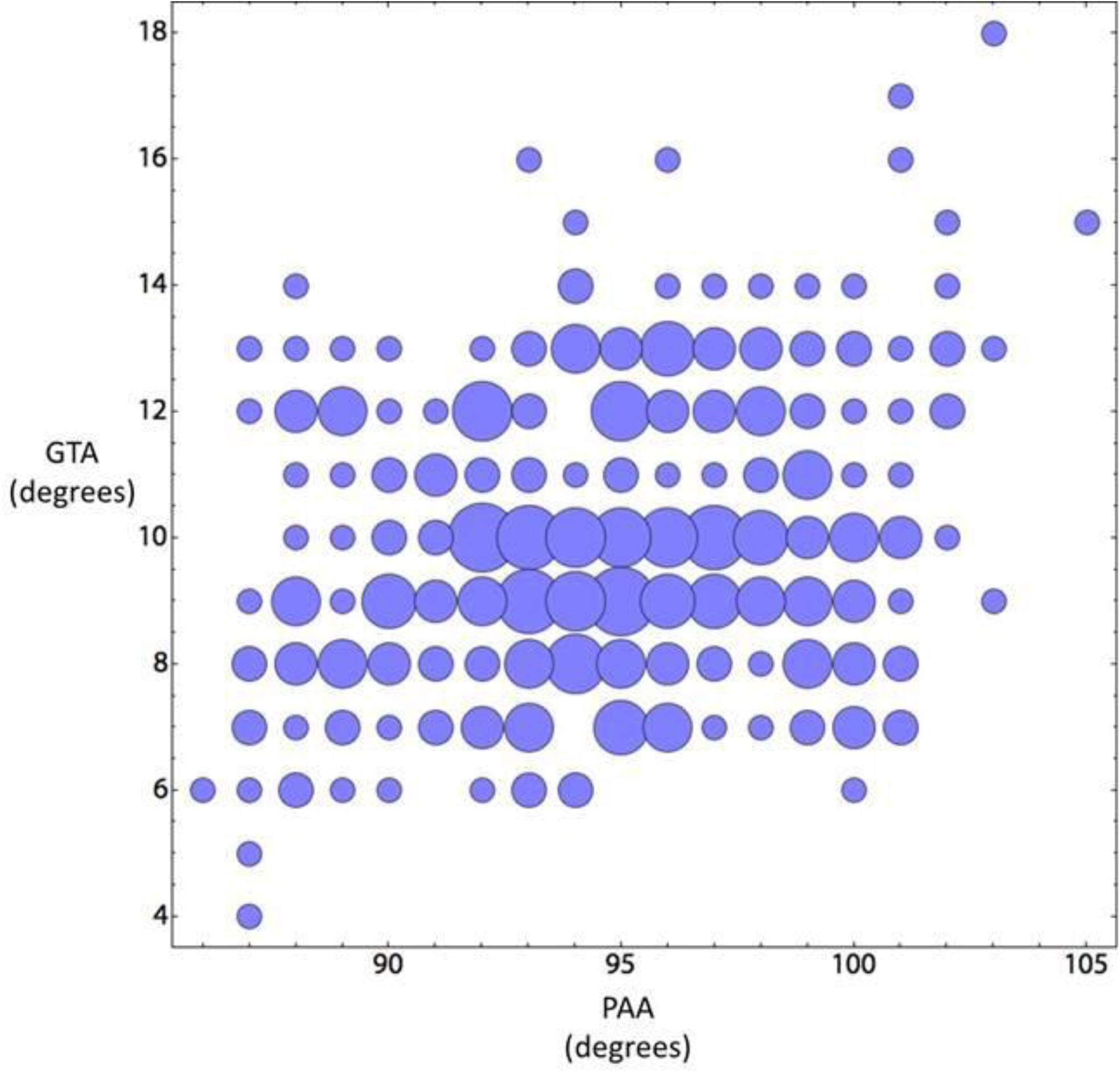
Relationship of first metatarsal proximal angle of articulation (PAA) and great toe angle (GTA) in living humans. Larger bubbles indicate more individuals of given values. Pearson’s r = 0.26 (p < 0.001); n = 331.

To test whether the two samples’ (living human and fossil hominin) PAA measurements are different enough that the difference in their means could be produced by chance, the resampling equivalent of a t-test (with no replacement due to the dramatic difference in sample sizes) was utilized. I asked if the known data universe produced by joining the two samples and choosing (without replacement) samples of the same sizes, then how often do those random samples have a mean difference that is greater than or equal to the difference in the means of the living human and fossil data? The result is an empirical p-value that tests whether or not the two samples are more different than expected by random chance. The p-value for the data after one million trials is 0.408; thus, the two sample means are not more different than expected by random chance. I also tested whether the two samples are more variable than expected by chance by examining the difference in standard deviation. The p-value for this test was very similar at 0.41.

## Discussion

Though the convergence of the hallux on the rest of the foot such that living humans are incapable of functional opposability but often retain limited functional grasping ability (compared to extant apes) is an important adaptation for bipedalism, the large sample of living humans in this study illustrates the impressive variability of this character found in modern populations. Furthermore, this sample is potentially biased *against* a full appreciation of hallucal convergence variability given that it is derived from a habitually shod population. That is, though the sample is highly variable, it is possible that a similarly large population of habitually unshod humans would exhibit even more variation, especially towards greater hallucal divergence.

The great variability of hallucal convergence present in this study’s sample of modern humans illustrates, succinctly, that there are many ways to assemble the legion soft-and hard tissues that each contribute to the relationship of the hallux with the foot in such a manner that permits unproblematic terrestrial bipedalism – while facilitating an ecologically important degree of arboreality. Though Latimer and Lovejoy (1990) argued “an opposable great toe is the *sine qua non* of primate arboreality,” I agree broadly with Susman *et al.* (1984) that hallucal abduction is not an essential condition of arboreality in hominins, but for different reasons than they argue. Venkataraman *et al.* (2013) poignantly observe that living humans have not entirely sacrificed arboreal mobility: their study of Twa hunter-gatherers vividly illustrates how modern human skeletons are capable of vertical climbing and arboreal resource exploitation, especially when complemented by soft-tissue anatomy well-accommodated to arboreal behavior. Though Venkataraman *et al*. (2013) emphasize the role of extraordinary ankle dorsiflexion in modern human exploitation of arboreal resources, they also highlight that many modern humans are capable of functional arboreal grasping with their halluces.

The results of this study imply the modern pattern of hallucal adduction, such that the great toe is capable of functional ‘clamping’ but not opposability is quite ancient. *Australopithecus* and early *Homo* exhibit a range of hallucal adduction variation encompassed by modern people; this implies their use of arboreal resources would have been no less than that of people who today exploit arboreal resources. That modern human hunter-gatherers often grasp arboreal substrates with their halluces (Fig. 4) and that even (nearly) exclusively terrestrial, habitually shod modern Midwestern Americans exhibit a range of variation that includes the hominin fossils surveyed in this study suggests long-term stabilizing selection maintaining foot and ankle anatomy that does not conform to the long-standing arboreal-terrestrial dichotomy that has dominated anatomical and behavioral interpretations of fossil hominins, consistent with the findings of Venkataraman *et al.* (2013).

**Figure 4.**
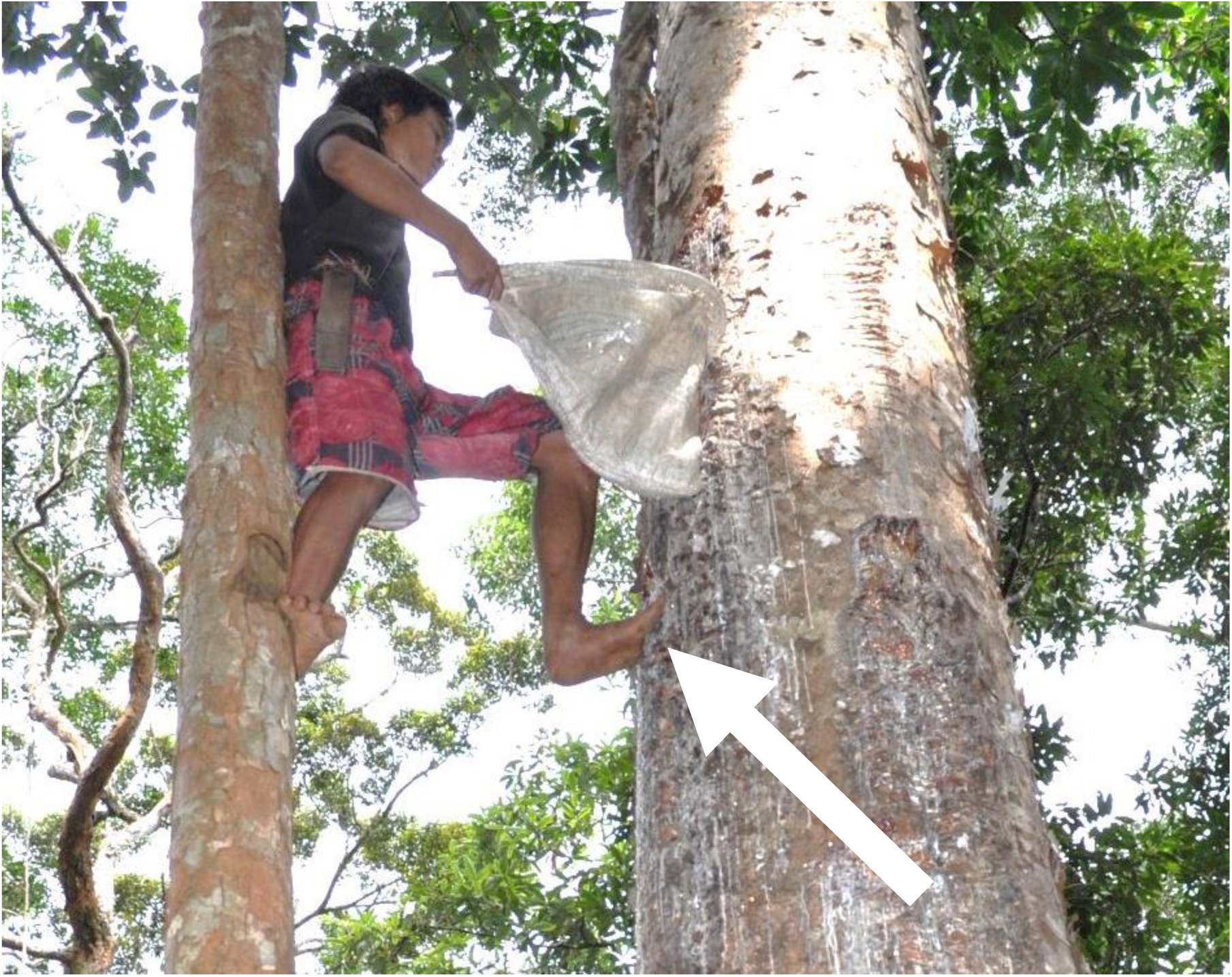
Climbing in modern humans. A Philippine Palawano man climbing a tree to collect gum. Note his usage of hallucal “clamping” on the bark of the tree. Photo courtesy Noah Theriault.

The presence of a fully abductable, opposable hallux limits but does not preclude terrestrial locomotion by apes. Similarly, the adducted, non-opposable hallux of modern humans limits but does not preclude our exploitation of arboreal resources. Venkataraman *et al.* (2013) caution that the range of modern human behavioral plasticity, specifically as it relates to arboreality, has been underappreciated by much previous research on the evolution of hominin foot and ankle anatomy as it reflects the evolution of hominin behavioral ecology. This study cautions that the range of modern human anatomical variability has also been underappreciated by much previous research on the evolution of hominin foot and ankle anatomy.

The anatomy of the BRT-VP-2/73 first metatarsal is different in several respects from the known foot bones of *A. afarensis,* and Haile-Selassie et al. (2012) noted its anatomical similarity to the foot of *Ardipithecus*. A broader comparison with a large sample of humans shows that BRT-VP-2/73 is consistent with the variation found in many modern humans. From a phylogenetic perspective, it may be premature to speculate that the BRT-VP-2/73 foot represents a taxon distinct from *A. afarensis*. If this foot specimen and the Hadar foot bones (and the Laetoli footprints) are representative of variation no wider than that found within living human populations, there is no reason to diagnose these remains as specifically distinct.

Biomechanically, I suggest that the Woranso-Mille foot’s second metatarsal diaphyseal medial torqueing is consistent with interphalangeal ‘clamping’ given its non-opposable hallux – though this notion will require further evaluation utilizing modern humans who regularly exploit arboreal resources by climbing. Whether there exists population-level differences between modern humans in variation of the first metatarsal PAA and the hallucal GTA remains to be determined. That said, I echo the qualification of Zipfel *et al*. (2009) in inferring too much from an isolated skeletal element, and caution that while the PAA of the first metatarsal is correlated with the relationship of the hallux with the foot, hallucal divergence can only be fully understood in a living, functional foot.

## Conclusion

This study highlights variability in the pattern of hallucal adduction within a population of modern, habitually shod humans. I suggest the proximal articulation angle of the first metatarsal provides evidence for the magnitude of hallucal convergence in fossil hominin foot remains that do not preserve the medial cuneiform. The correlation of the first metatarsal’s proximal articulation angle with the great toe angle derived from living humans with their soft tissues intact can also provide information about hallucal adduction in more complete hominin fossil feet. I find that as in modern humans, fossil hominins exhibit a remarkable degree of variation in hallucal adduction. How populations of hominins varied compared to modern humans will require additional fossil material, and identification of additional skeletal correlates of hallucal adduction.

